# Liposomal iron bisglycinate hydrochloride, but not free iron bisglycinate, enhances serum iron restoration in a mouse model of inflammation-induced anemia

**DOI:** 10.1101/2025.07.28.667108

**Authors:** Meygal Kahana Sela, Amir Agbaria, Noor Omari Knani, Rami Awida, Abraham Weizman, Moshe Gavish

## Abstract

**Background/Objectives:** Iron deficiency is a major global health concern, particularly in inflammation-related anemia (IRA), where iron absorption and mobilization are impaired. Liposomal encapsulation offers a promising strategy to enhance the bioavailability and efficacy of iron supplements. This study aimed to evaluate the therapeutic efficacy of orally administered liposomal iron bisglycinate hydrochloride (LIBH), compared to free iron bisglycinate hydrochloride (FIBH), in restoring serum iron levels in a mouse model of lipopolysaccharide (LPS)-induced anemia of inflammation (LIAI).

**Methods:** Male C57BL/6 mice received intraperitoneal LPS (5 mg/kg) to induce LIAI and were simultaneously treated with either FIBH or LIBH (1 mg/kg, oral). Liposomes (150 - 200 nm) exhibited high encapsulation efficiency (93%) and stability (−38 to -45 mV zeta potential). Serum iron levels were measured 24 hours post-treatment.

**Results:** LPS administration significantly reduced serum iron levels. LIBH restored serum iron levels by 35 - 65% compared to baseline (p < 0.005), outperforming FIBH (0 - 16%, p = NS). Electron microscopy confirmed the structural integrity of LIBH liposomes.

**Conclusion:** LIBH supplementation significantly improves serum iron levels in LIAI and may represent a superior alternative to traditional free iron therapy, particularly in inflammatory conditions. Further studies are warranted to assess its efficacy in other models of anemia.

## Introduction

Iron is critical in oxygen transport, neurotransmitter synthesis, DNA replication, and energy metabolism. Maintaining iron homeostasis is essential, and disruptions can lead to significant health issues, including iron deficiency anemia (IDA). The World Health Organization estimates that up to two billion people worldwide suffer from IDA [1]. Oral iron supplementation remains the mainstay treatment for IDA, with commonly used formulations including ferrous sulfate, ferrous gluconate, and ferrous fumarate [2,3]. However, these compounds often cause gastrointestinal side effects such as nausea, constipation, and diarrhea, and have relatively poor bioavailability, typically less than 20% [4,5]. This necessitates high daily doses and often results in poor patient compliance. Anemia of inflammation (AI), also called anemia of chronic disease, is the most prevalent form of anemia among hospitalized patients. It is characterized by low serum iron and increased iron stores due to impaired iron absorption and mobilization driven by elevated hepcidin and inflammatory cytokines [6]. Traditional iron supplementation is often ineffective in AI.

Ferrous bisglycinate, a chelated form of iron, has demonstrated superior bioavailability and a better side-effect profile than conventional salts [7]. Liposomes, spherical vesicles composed of phospholipid bilayers, are effective drug carriers due to their stability, biocompatibility, and ability to enhance oral bioavailability [8,9].

We used a lipopolysaccharide (LPS)-induced mouse model of AI to evaluate whether liposomal iron bisglycinate hydrochloride (LIBH) could improve serum iron levels more effectively than free iron bisglycinate hydrochloride (FIBH). We hypothesized that the enhanced delivery properties of LIBH would result in superior correction of inflammation-induced hypoferremia.

## Materials and Methods

### Liposome preparation

Liposomes were prepared using dioleoylphosphatidylcholine/ cholesterol /alpha-lipoic acid (DOPC/Chol/ALA) in a molar ratio of 5.5:4.0:0.5. DOPC, Chol and ALA were dissolved in ethanol. Solvents were allowed to evaporate using rotary evaporation at 37°C. The dried lipid film was hydrated with either double-distilled water (DDW) for empty liposomes or a DDW solution of iron bisglycinate hydrochloride for LIBH. The mixture was sonicated and subjected to six freeze-thaw cycles. Liposomes were extruded through filters of decreasing pore size (400 nm, 200 nm, 100 nm) and dialyzed to remove unencapsulated iron.

The resulting liposomes had a uniform 150–200 nm size, a zeta potential of −38 to −45 mV, and an iron encapsulation efficiency of 93%. Stability was confirmed for two weeks at 4°C.

Male C57BL/6 mice (9–10 weeks, 25 ± 2 g) were housed under standard conditions and approved protocols. The study was approved by the Technion’s Committee for Experiments in Animals (IIT - IL - 2507 - 177).

Mice were divided into six groups:

1. Naïve: Untreated
2. Vehicle: Empty liposomes (50 µL, oral)
3. LPS: LPS (5 mg/kg, I.P.)
4. LPS + FIBH: LPS + free iron bisglycinate (1 mg/kg, 50 µL, oral)
5. LPS + ∼700nm LIBH: LPS + liposomal iron bisglycinate **without** extruder (1 mg/kg, 50 µL, oral)
6. LPS + ∼150nm LIBH: LPS + liposomal iron bisglycinate **with** extruder (1 mg/kg, 50 µL, oral)

### Blood Collection and Serum Iron Measurement

Twenty-four hours post-treatment, mice were euthanized, and blood was collected. Serum was separated by centrifugation and stored at 4°C till analysis. Serum iron levels were measured using the Architect ci16200 analyzer (Abbott Core Laboratory, Abbott Park, IL, USA).

### Electron Microscopy

Liposome morphology and size were assessed using transmission electron microscopy. Liposomes were stained with uranyl acetate and imaged with a Talos L120C microscope (FEI Israel Thermofisher, Airport City, Israel).

## Results

### Characterization of the liposomes

#### Liposomal and free iron preparation efficacy for serum iron levels in mice

The serum iron levels of C57BL/6 mice were assessed 24 h after administering LPS (I.P., 5 mg/kg) with or without oral co-administration of iron (0.93 mg/kg). In the Serum iron levels decreased considerably 24 h after LPS administration. In a mouse model of LPS-induced anemia of inflammation (LIAI) the serum iron level decreased by 67% (*p* = 0.002), while co-administration with FIBH did not affect the serum iron level (68%; *p* < 0.0001), and co-administration with LIBH without extruder use (□700nm) resulted in an significantly the serum iron level (17%; *p* = 0.417). In contrast, administration of LIBH modified by extruder use (□150nm; LPS+∼150nm LIBH) increased the serum iron level by 35% compared to LPS alone (*p* = 0.0059) and by 31% compared to the LPS+FIBH group (*p* = 0.036) and 22% compared to the LPS+∼700nm LIBH subgroup.

To validate the results, we repeated the experiment with larger groups and without the LPS+ 700nm LIBH group. As shown in Figure 2, similar to the original experiment (Figure 1), serum iron levels decreased significantly 24 h after LPS administration. In the LPS group, the serum iron level decreased by 64% (*p* < 0.0001), while co-administration with FIBH did not affect the serum iron level (10%; *p* = 0.40). In contrast, treatment with LIBH modified by extruder use (□ 150nm; LPS+ □ □150nm LIBH) increased the serum iron level by 55% compared to LPS alone (*p* < 0.0001) and by 50% compared to FIBH (*p* < 0.0001).

**Figure 1.**
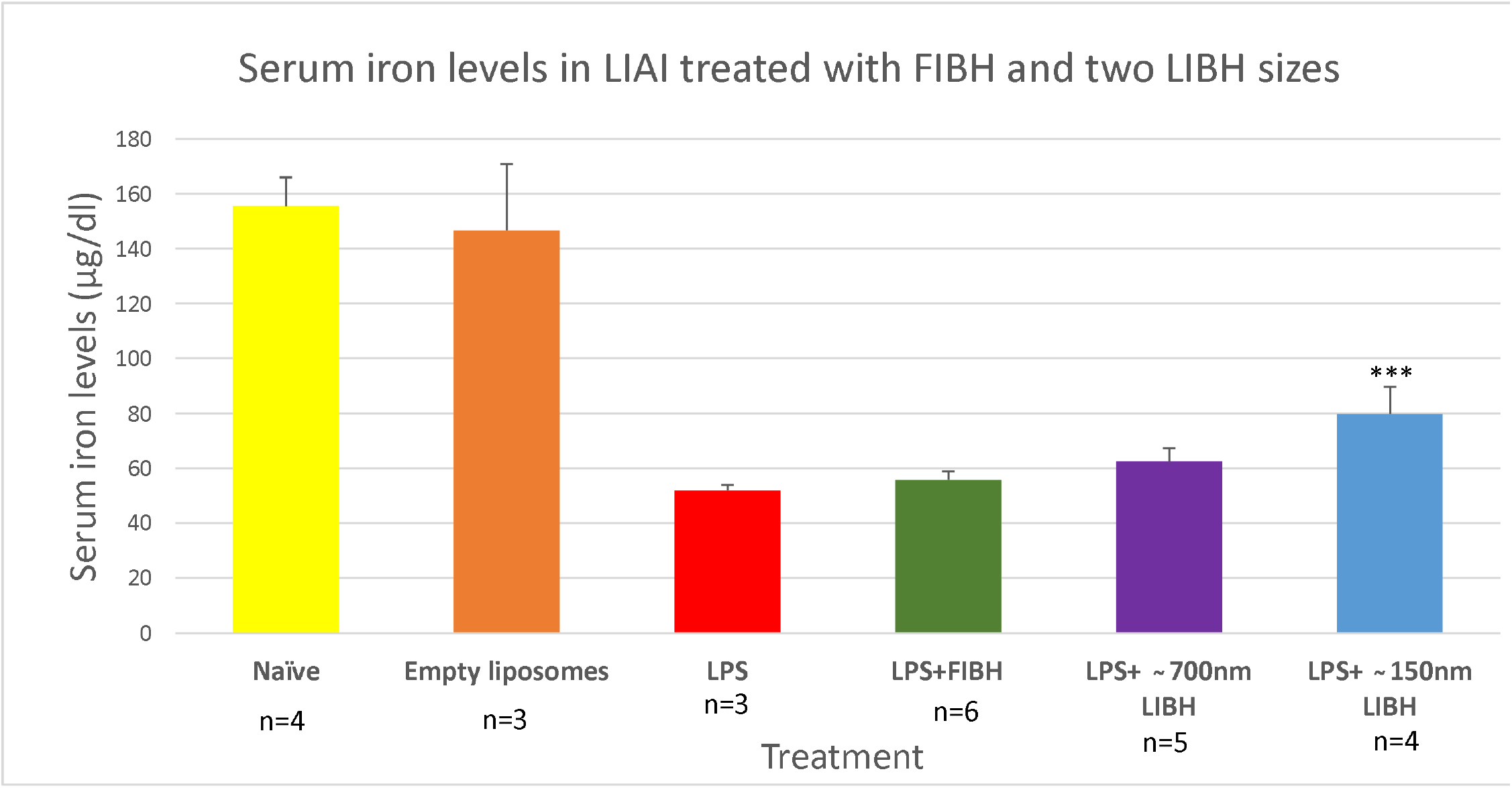
The impact of free iron bisglycinate hydrochloride (FIBH) and two sizes (∼150nm, ∼700nm) of liposomal iron bisglycinate hydrochloride (LIBH) on serum iron levels in a mouse model of lipopolysaccharide (LPS)-induced anemia of inflammation (LIAI). One-way ANOVA: F =19.66 (DF 5, 19), p <0.0001. LPS+ □ □150nm LIBH: p=0.036 versus LPS+FIBH and p= 0.0059 versus LPS alone. LPS+□ □ 700nm LIBH did not differ significantly from LPS or LPS+FIBH.

**Figure 2.**
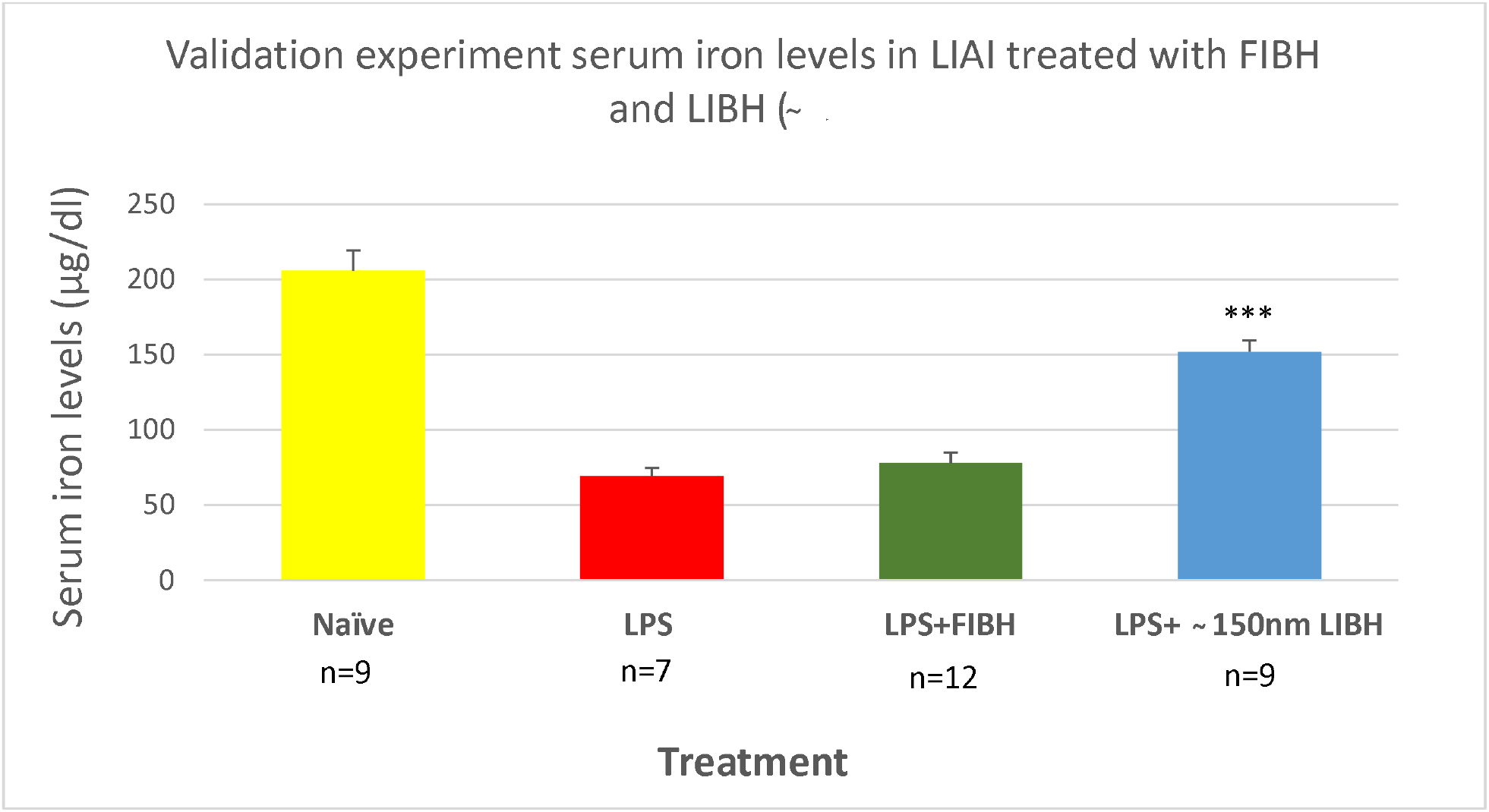
Serum iron levels in lipopolysaccharide (LPS)-induced anemia of inflammation (LIAI) mice treated with free iron bisglycinate hydrochloride (FIBH) and liposomal iron bisglycinate hydrochloride (LIBH) (□ □150nm). One-way ANOVA: F =49.03 (DF 3, 33), p <0.0001. LPS+ □ □150nm LIBH p<0.0001: versus LPS+FIBH, p< 0.0001 versus LPS.

#### ∼150nm LIBH Stability

Assessments were performed over 35 days (n=3 at each time point) to evaluate the stability of the ∼150nm LIBH characteristics. Table 1 shows the stability of the liposome characteristics.

**Table 1:** ∼150nm LIBH stability over 35 days

| Day | Size (nm) | Zeta potential (mV) | Polydispersity Index |
| --- | --- | --- | --- |
| 0 | 175 | -35.3 | 0.06 |
| 7 | 160 | -36.7 | 0.04 |
| 28 | 139 | -35.5 | 0.1 |
| 35 | 181 | -29 | 0.1 |

**Legend:** LIBH - liposomal iron bisglycinate hydrochloride

#### Electron microscopy of the liposomes

Electron microscopy (EM) shows a representative homogeneous liposome of size 150 nm and the absence of aggregation or multilayer (Figure 3).

**Figure 3.**
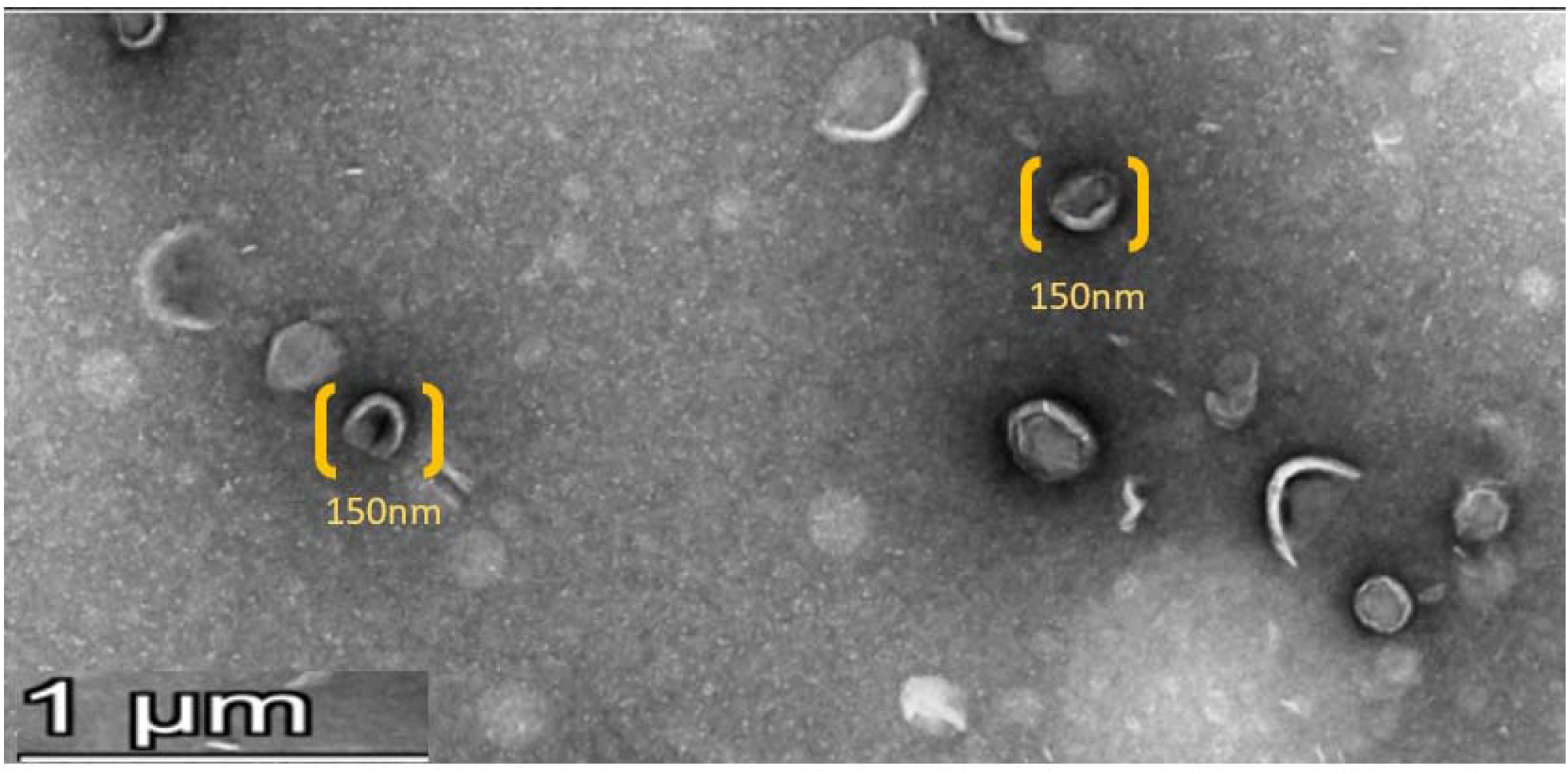
Size assessment of the ∼150nm liposomes using electron microscopy A representative electron microscopy of liposomes showing them to be intact (no aggregation or multilayer) and of a stable size of ∼150 nm.

## Discussion

We demonstrated that oral ∼150nm LIBH significantly improves serum iron restoration in an LPS-induced mouse model of inflammation-related anemia, outperforming FIBH. Liposomal encapsulation of iron facilitates absorption, may protect against gastrointestinal irritation, and is a safe and tolerable delivery system [10-12]. Our ∼150nm LIBH formulation was physically stable, with consistent size and zeta potential over 35 days. The observed zeta potential (> |30| mV) suggests good stability. Moreover, the polydispersity index values of 0.1 or less, indicate homogeneity of the liposomes, which is crucial for reproducible oral delivery.

In contrast to FIBH, LIBH is shielded from oxidative degradation and the generation of reactive oxygen species. This protective encapsulation helps preserve the integrity and bioactivity of iron during gastrointestinal transport and systemic circulation [9,13,14]. Additionally, by minimizing direct contact with the intestinal mucosa, liposomal delivery reduces the risk of local irritation and gastrointestinal side effects (e.g., diarrhea, constipation, and dyspepsia) that reduce adherence to oral iron administration [15-17] and are commonly associated with FIBH supplementation. Our findings suggest that LIBH could present a valuable alternative for patients with inflammation-related anemia (IRA) and potentially other iron deficiency states. Further studies in humans are necessary to confirm these promising preclinical results.

## Author Contributions

Conceptualization, M.K.S., A.A., N.O.K., R.A., A.W. and M.G.; Methodology, M.K.S, A.A., N.O.K., A.W. and M.G.; Formal Analysis, M.K.S., A.A. and N.O.K.; Investigation, M.K.S., A.A. and N.O.K.; Resources, R.A. and M.G.; Data Curation, M.K.S., A.A. and N.O.K.; Writing – Original Draft Preparation, M.K.S., A.A., N.O.K. and A.W.; Writing – Review &Editing, A.W. and M.G.; Visualization, M.K.S., A.A. and N.O.K.; Supervision, A.W. and M.G.; Project Administration, M.K.S., A.A. and N.O.K.; Funding Acquisition, R.A.”

## Institutional Review Board Statement

The study was conducted according to the guidelines of the Declaration of Helsinki, and approved by the Institutional Review Board (or Ethics Committee) of the Technion - Israel Institute of Technology (protocol code IIT - IL - 2507 - 177, July 2025).

## Data Availability Statement

Dataset available on request from the authors.

## References

1. Armitage, A.E.; Drakesmith, H. Genetics. The battle for iron. Science 2014, 346, 1299–1300, doi:10.1126/science.aaa2468.

2. Wilson, K.; Sloan, J.M. Iron-Deficiency Anemia. N Engl J Med 2015, 373, 485, doi:10.1056/NEJMc1507104.

3. Cancelo-Hidalgo, M.J;. Castelo-Branco, C.; Palacios, S.; Haya-Palazuelos, J.; Ciria-Recasens, M.; Manasanch, J.; Pérez-Edo, L. Tolerability of different oral iron supplements: a systematic review. Curr Med Res Opin 2013, 29, 291–303, doi:10.1185/03007995.2012.761599.

4. Patil, P.; Geevarghese, P.; Khaire, P.; Joshi, T.; Suryawanshi, A.; Mundada, S.; Pawar, S.; Farookh, A. Comparison of Therapeutic Efficacy of Ferrous Ascorbate and Iron Polymaltose Complex in Iron Deficiency Anemia in Children: A Randomized Controlled Trial. Indian J Pediatr 2019, 86, 1112–1117, doi:10.1007/s12098-019-03068-2.

5. Tondeur, M.C.; Schauer, C.S.; Christofides, A.L.; Asante, K.P.; Newton, S.; Serfass, R.E.; Zlotkin, S.H. Determination of iron absorption from intrinsically labeled microencapsulated ferrous fumarate (sprinkles) in infants with different iron and hematologic status by using a dual-stable-isotope method. Am J Clin Nutr 2004, 80, 1436–1444, doi:10.1093/ajcn/80.5.1436.

6. Madu, A.J.; Ughasoro, M.D. Anaemia of Chronic Disease: An In-Depth Review. Med Princ Pract 2017, 26, 1–9, doi:10.1159/000452104.

7. Fischer, J.A.J.; Cherian, A.M.; Bone, J.N.; Karakochuk, C.D. The effects of oral ferrous bisglycinate supplementation on hemoglobin and ferritin concentrations in adults and children: a systematic review and meta-analysis of randomized controlled trials. Nutr Rev 2023, 81, 904–920, doi:10.1093/nutrit/nuac106.

8. Yuba, E.; Harada, A.; Sakanishi, Y.; Watarai, S.; Kono, K. A liposome-based antigen delivery system using pH-sensitive fusogenic polymers for cancer immunotherapy. Biomaterials 2013, 34, 3042–3052, doi:10.1016/j.biomaterials.2012.12.031.

9. Xu, Z.; Liu, S.; Wang, H.; Gao, G.; Yu, P.; Chang, Y. Encapsulation of iron in liposomes significantly improved the efficiency of iron supplementation in strenuously exercised rats. Biol Trace Elem Res 2014, 162, 181–188, doi:10.1007/s12011-014-0143-0.

10. Riva, A.; Ronchi, M.; Petrangolini, G.; Bosisio, S.; Allegrini, P. Improved Oral Absorption of Quercetin from Quercetin Phytosome®, a New Delivery System Based on Food Grade Lecithin. Eur J Drug Metab Pharmacokinet 2019, 44, 169–177, doi:10.1007/s13318-018-0517-3.

11. Iwanaga, K.; Ono, S.; Narioka, K.; Kakemi, M.; Morimoto, K.; Yamashita, S.; Namba, Y.; Oku, N. Application of surface-coated liposomes for oral delivery of peptide: effects of coating the liposome’s surface on the GI transit of insulin. J Pharm Sci 1999, 88, 248–252, doi:10.1021/js980235x.

12. Chirra, H.D.; Desai, T.A. Emerging microtechnologies for the development of oral drug delivery devices. Adv Drug Deliv Rev 2012, 64, 1569–1578, doi:10.1016/j.addr.2012.08.013.

13. Niu, M.; Lu, Y.; Hovgaard, L.; Guan, P.; Tan, Y.; Lian, R.; Qi, J.; Wu, W. Hypoglycemic activity and oral bioavailability of insulin-loaded liposomes containing bile salts in rats: the effect of cholate type, particle size and administered dose. Eur J Pharm Biopharm 2012, 81, 265–272, doi:10.1016/j.ejpb.2012.02.009.

14. Kaddah, S.; Khreich, N.; Kaddah, F.; Charcosset, C.; Greige-Gerges, H. Cholesterol modulates the liposome membrane fluidity and permeability for a hydrophilic molecule. Food Chem Toxicol 2018, 113, 40–48, doi:10.1016/j.fct.2018.01.017.

15. Patel, H.M.; Ryman, B.E. Oral administration of insulin by encapsulation within liposomes. FEBS Lett 1976, 62, 6, 0-63doi:10.1016/0014-5793(76)80016-6.

16. Gradauer, K.; Barthelmes, J.; Vonach, C.; Almer, G.; Mangge, H.; Teubl, B.; Roblegg, E.; Dünnhaupt, S.; Fröhlich, E.; Bernkop-Schnürch, A., et al. Liposomes coated with thiolated chitosan enhance oral peptide delivery to rats. J Control Release 2013, 172, 872–878, doi:10.1016/j.jconrel.2013.10.011.

17. Zariwala, M.G.; Bendre, H.; Markiv, A.; Farnaud, S.; Renshaw, D.; Taylor, K.M.; Somavarapu, S. Hydrophobically modified chitosan nanoliposomes for intestinal drug delivery. Int J Nanomedicine 2018, 13, 5837–5848, doi:10.2147/ijn.S166901.

